# Assessing the Relationship Between Vegetation Structure and Harvestmen Assemblage in an Amazonian Upland Forest

**DOI:** 10.1101/078220

**Authors:** Pío Colmenares, Fabrício B. Baccaro, Ana Lúcia Tourinho

**Author notes:** Corresponding author: Ana Lúcia Tourinho.

## Abstract

1. Arthropod diversity and non-flying arthropod food web are strongly influenced by habitat components related to plant architecture and habitat structural complexity. However, we still poorly understand the relationship between arthropod diversity and the vegetation structure at different spatial scales. Here, we examined how harvestmen assemblages are distributed across six local scale habitats (trees, dead trunks, palms, bushes, herbs and litter), and along three proxies of vegetation structure (number of palms, number of trees and litter depth) at mesoscale.

2. We collected harvestmen using cryptic manual search in 30 permanent plots of 250 m at Reserva Ducke, Amazonas, Brazil. The 30 plots cover approximately 25 km^2^ of upland forests. At a local scale, harvestmen were most diverse and abundant on trees. The likely preference of trees by harvestmen may be related to the variety of local microhabitats offered by large trees. However, despite the strong link between number of harvestman species and individuals with large trees, only harvestmen assemblages composition were related with number of trees and with number of palms, at mesoscale.

3. Harvestman richness and abundance were not related with any vegetation structure predictor at mesoscale. Therefore, areas of *upland* forest in the central Amazon with large trees and palms do not harbor more harvestman species nor individuals, but are suitable to maintain different harvestmen assemblages.

## Introduction

Tropical forests occupy 11% of the earth’s surface yet maintain more than 60% of its terrestrial biodiversity (Wilson, 2000). The reason that promotes such highly concentration of plant and animal diversity remains contentious (Hubbell, 2001; Novotny *et al*., 2006), but there is compelling evidence that tropical diversity is influenced and maintained by wide environmental gradients and habitat structural heterogeneity (Gardner *et al*., 1995; Halaj *et al*., 2000). In the Amazon rainforest several arthropod assemblages have been characterized and related to environmental gradients. For instance, ant diversity is affected by water table depth variation (Baccaro *et al*., 2013), ants influence termites (Pequeno & Pantoja, 2012), different environmental predictors affect cockroaches simultaneously (Tarli *et al*., 2013), and understory forest structure affect harvestmen assemblages (Tourinho *et al*., 2014; Porto *et al*., 2016). However, few studies investigated how non-flying arthropods are limited or affected by vegetation structure (Vasconcelos *et al*., 2008; Donoso *et al*., 2010) and we still poorly understand the role of vegetation structure on arthropod diversity in forested areas at different spatial scales.

Arachnids represent one of the most diverse groups of arthropods, and around 2% of described species occurs in the Amazon basin (Adis & Harvey, 2000). With more than 6600 species described (Kury, 2016), harvestmen represent the third most diverse order of arachnids, after spiders and mites, and are well represented in the Amazon biome (Kury & Pinto-da-Rocha, 2002). Harvestmen are mostly predators and strongly affected by temperature and humidity, thus, susceptible to dehydration (Curtis & Machado, 2007).

Only a few studies have investigated the relationship between harvestmen species and habitat structure. It was demonstrated that harvestmen assemblages from Atlantic forest are positive related to forest quality, responding more drastically to fragmentation than most arthropods (Bragagnolo *et. al.*, 2007). Proud et al., (2012) suggests that in tropical forests of Costa Rica, harvestmen can use trees as refuges when disturbed, but only in sites with higher harvestmen diversity in the ground/litter microhabitat. Recent evaluation of collecting techniques also offered evidence for a relationship between number of palm trees and harvestmen assemblage composition in upland Amazonian forest (Tourinho *et al*., 2014; Porto *et al*., 2016). However, how vegetation structure directly or indirectly influences harvestmen assemblages remains still little understood.

Here, we investigate the relationship between vegetation structure and a harvestmen assemblage in an *upland* forest in the Central Amazon at two different spatial scales. We investigate how harvestmen assemblages are distributed at local scale (plots of 500 m^2^ each). We also test and describe the relationship of two direct proxies of vegetation structure (number of palms and number of trees) and one indirect proxy of vegetation structure (litter depth) with the harvestmen assemblage composition at mesoscale (25 km^2^).

## Methods

### Study area

The study area is located in the central Amazon, in the Reserva Forestal Adolpho Ducke (Fig. 1), which is a 100 km^2^ fragment of *terra firme* forest administrated and protected by the Instituto Nacional de Pesquisas da Amazônia (INPA). The vegetation is typically upland rainforest, with a diversity of trees around 1200 species (Costa et al., 2009), with a canopy height of 30–35m above the ground. Annual mean temperature is 26ºC. Annual precipitation is between 1.900–2.300 mm^3^, and the wet season usually begins in November and lasts until May (Baccaro et al., 2008). Altitudinal variation is between 30–180 m asl.

**Fig. 1.**
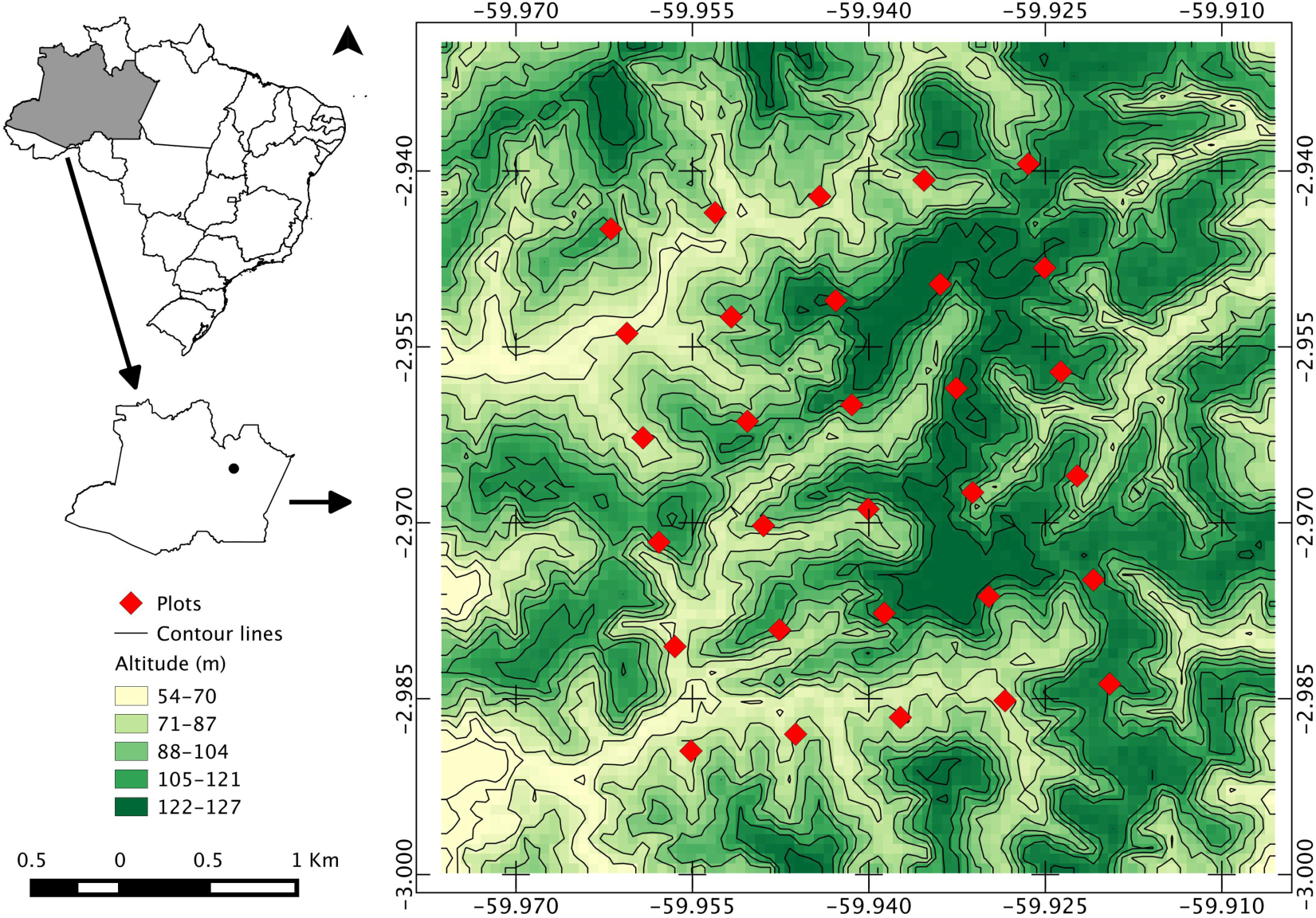
Relative position of the 30 sampled plots at Reserva Forestal Adolpho Ducke.

### Sampling design

A total of 30 plots were sampled between July and November 2014, covering an area of 25 km^2^ (Fig. 1). Collecting plots were established following the RAPELD protocol (Magnusson *et al*., 2005). Each plot is a 250 m transect with variable width, and the distance between them is 1 km. Two experienced collectors sampled every plot in a single visit for one hour. The sampling were undertaken along one meter to each side of the main line of the 250 m-long plots, totalizing 500 m^2^ of sampled area per plot (2 x 250 m). Along the survey, all harvestman found within the plot were collected. The habitat occupied by each individual was recorded at the moment of the capture, and were classified into trees, dead trunks, palms, bushes, herbs and litter. Harvestmen were collected using cryptic manual searching, which focus on specific habitats described above (Porto *et al*., 2016). This method allows for collection of more species and individuals compared with traditional surveys (Porto *et al*., 2016).

### Species data

To identify harvestmen species we examined the external morphology under a stereomicroscope and compared with the original descriptions provided in the literature (Pinto-da-Rocha, 1994, 1996, 1997, 2004; Kury, 2003) type material or pictures of the type material. Nymphs and females with ambiguous morphology were excluded. In the case of groups with a very conservative external morphology and/or poorly understood taxonomy (e.g. Cosmetidae, Sclerosomatidae, Zalmoxidae) we also prepared their male genitalia to allow proper species delimitation, following Acosta et al., (2007). Material is labeled and deposited in the Invertebrate collection at INPA (curator Celio Magalhães).

### Environmental data

We tested two direct proxies of vegetation structure as habitats available for harvestmen: number of palms (NPalm) and number of trees with diameter at breast height > 30 cm (DBH30); and litter depth (Litter) as an indirect proxy of vegetation structure. The number of trees, palms and litter depth were obtained from the data repository of the PPBio program (ppbio.inpa.gov.br). Within plots, all trees with diameter at breast height > 30 cm (DBH30) in 1 ha (40 m x 250 m) were mapped, and their diameters at 1.30 m (DBH) measured (Castilho et al., 2006). The same protocol was used to count and map the palm trees within plots. At every 5 m along the long axis of the plots, a measurement of litter depth was taken. Measurements consisted of forcing a stick of 0.5 cm in diameter into the litter until it reached the soil and noting the distance in cm between the top piece of litter and the soil. In addition, we also measured the diameter at breast high of trees with harvestmen during the sampling surveys.

### Data analysis

We generated two data matrices: one for the local scale analysis, using the habitats recorded at the moment of the capture (trees, dead trunks, palms, bushes, herbs and litter) as objects and species as columns, and another matrix for the mesoscale analysis using plots as objects and species as columns.

For species richness we used the total number of species collected per plot, for abundance we used the total number of individuals sampled per plot. We compared the number of species and abundance per plot (local scale) between each habitat predictor using analysis of variance ANOVA. Tukey´s Honest Significance Difference method was used to compute the 95% confidence interval for each factor. Residual analyses were used to investigate model assumptions.

For the mesoscale analysis, species composition per plot was summarized in a single multivariate axis using Non-metric Multidimensional Scaling NMDS, based on Bray-Curtis dissimilarity distances of the original abundance matrix. To evaluate the effect of vegetation structure on the harvestmen assemblage, we performed multiple regressions for each dependent variables against the independent variables, as follows: NMDS axis = *a*+*b*(Palms)+*b*(DBH30)+*b*(Litter), richness = *a*+*b*(Palms)+*b*(DBH30)+*b*(Litter), and abundance = *a*+*b*(Palms)+*b*(DBH30)+*b*(Litter). Partial regression plots were generated to show the relationships between variables. All three independent variables showed low correlation (r < 0.3).

## Results

A total of 689 adult harvestmen were collected, representing 27 species and 12 families (Table 1). The most abundant families were Cosmetidae (37.44% of total abundance) and Sclerosomatidae (22.78% of total abundance). The most common species were *Eucynortella duapunctata* (183 individuals), *Caluga* sp. 1 (83 individuals) and *Cynorta* sp. 1 (75 individuals). Abundance per plot ranged from nine to 54, with a mean of 22.96 individuals. Richness per plot varied between four to 13, with a mean of 7.93 species.

**Table 1.**
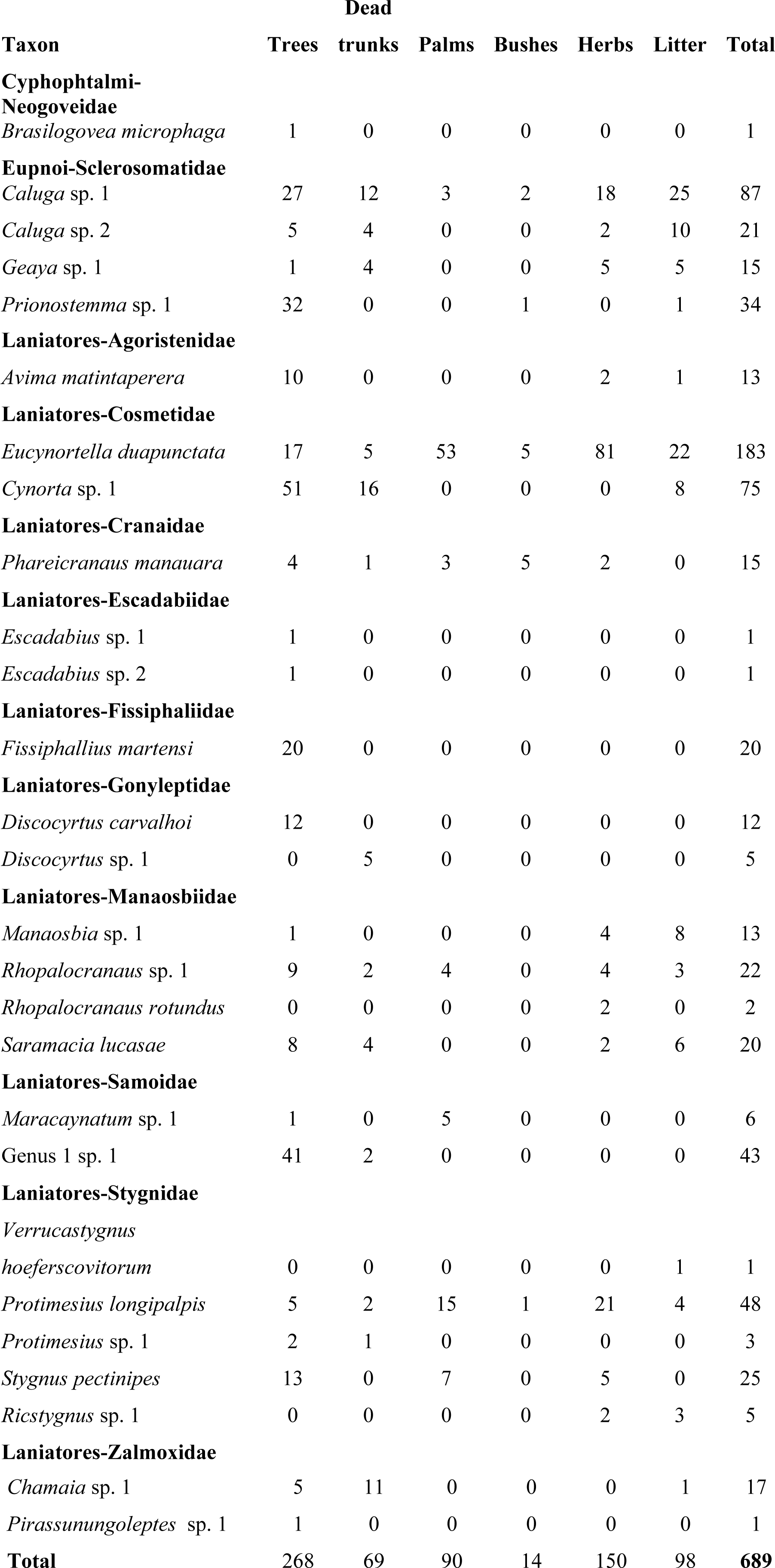
Species collected at Reserva Ducke, and the habitats they occupied

With 85.18% of species and 38.89% of individuals sampled, trees harbored the most diverse and abundant harvestman assemblage (ANOVA F_5,174_ = 15.16, P < 0.001 and F_5,175_ = 55.13, P < 0.001 respectively) (Figs. 2–3). The second most diverse habitat was herbs, while the third most abundant habitat was litter, but no strong effects were detected among other habitats. Bushes were the less diverse and harbors fewer harvestmen. Richness and abundance for the six habitats evaluated are summarized in table 2.

**Fig. 2.**
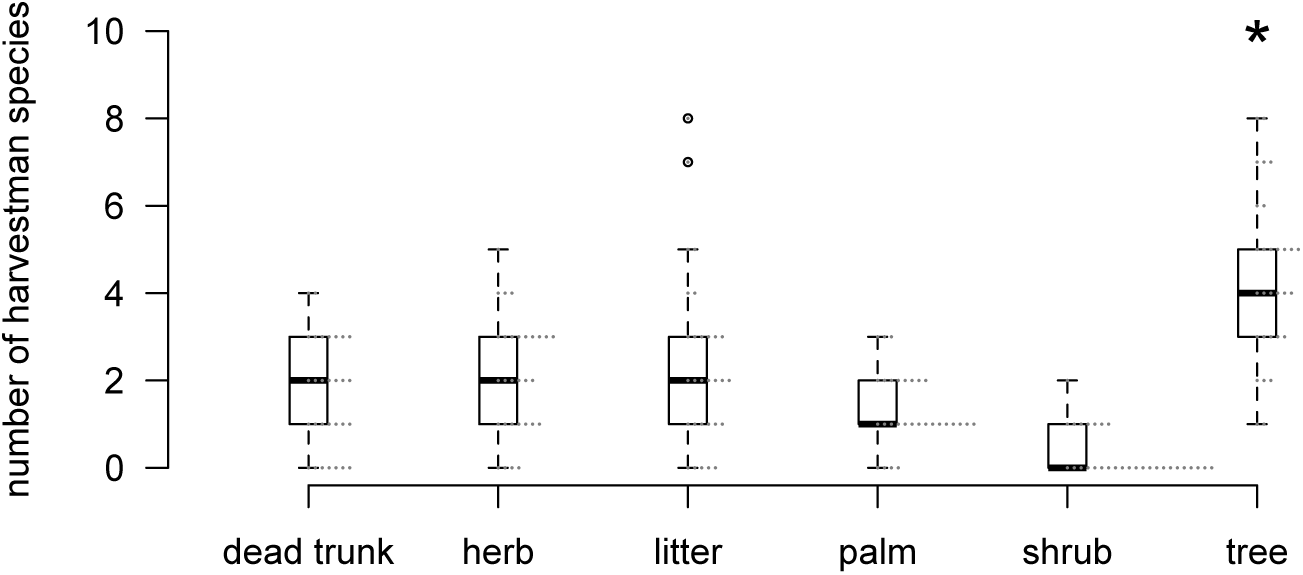
Number of harvestmen species sampled per habitat. Asterisk indicates significative differences.

**Fig. 3.**
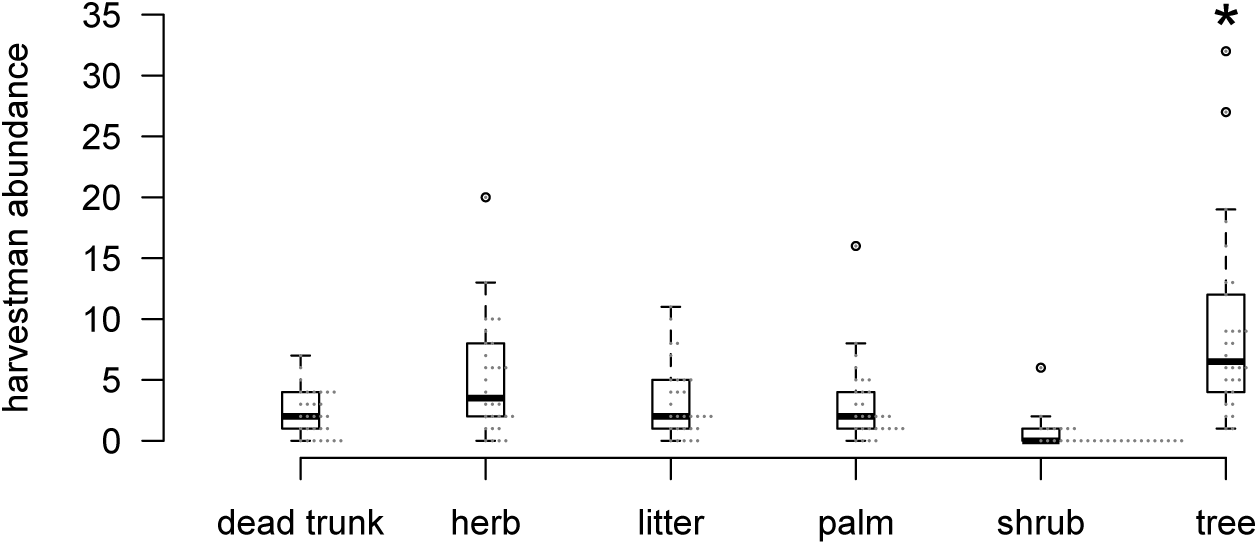
Abundance of harvestmen sampled per habitat. Asterisk indicates significative differences.

**Table 2.**
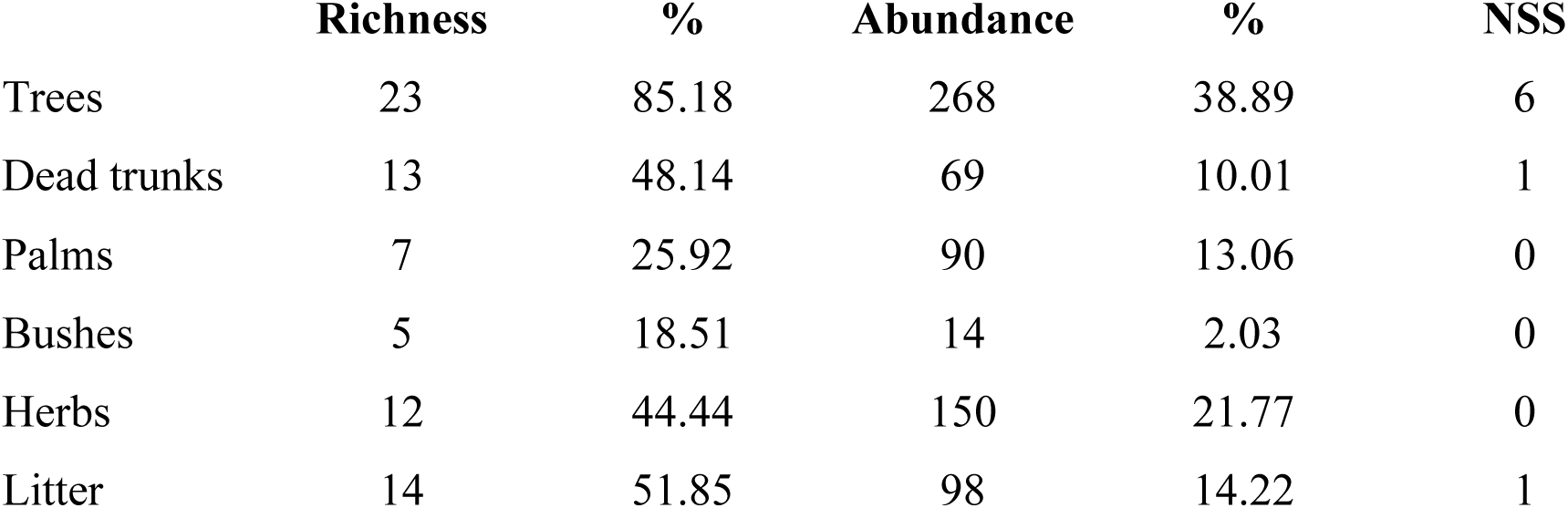
Distribution of harvestmen across sampled habitat at Reserva Ducke. NSS = number of species sampled only on a given habitat

Smaller Stygnidae such as *Verrucastygnus hoeferscovitorum* and *Ricstygnus* sp. 1 showed preference for the litter habitat, occasionally going on short herbs. Gonyleptids species of *Discocyrtus* showed habitat segregation, with individuals of *Discocyrtus carvalhoi* occurring only on live trees, and individuals of *Discocyrtus* sp. 1 occurring on fallen dead trunks. The two Escadabiidae collected were found on large trees with corrugated bark. All individuals of the Fissiphalliidae collected in our study, *Fissiphallius martensi*, were detected on trees with corrugated bark. As something unusual, we found one specimen of *Brasilogovea microphaga* climbing on a tree, which represents the first record of climbing behavior for a Cyphophtalmi.

The number of trees with diameter at breast height > 30 cm varied between 87 to 128 per plot (mean = 105.2). Conversely, the number of palms showed a wider range, varying between 97 to 448 palm trees per plot (mean = 269.7). The litter depth also varied largely between plots, ranging from only 1.31 to 4.18 cm (mean = 2.3 cm).

The NMDS ordination axis captured 59.55% of the variation of the species composition data (F = 640.4; DF = 1,433; P < 0.001). The multiple regression model, with the species composition as the response variable (NMDS Axis), explained 41.7% of the variation in the data (r^2^ = 0.417, P = 0.002) (Fig. 4). The independent variables that contributed significantly to the model were number of palms (b = 0.450, P = 0.005) and number of trees with diameter above breast height > 30 cm (b = 0.346, P = 0.036). Litter depth did not affect species composition. Multiple regression models for richness and abundance were non-significant. Results of regression models are summarized in table 3.

**Fig. 4.**
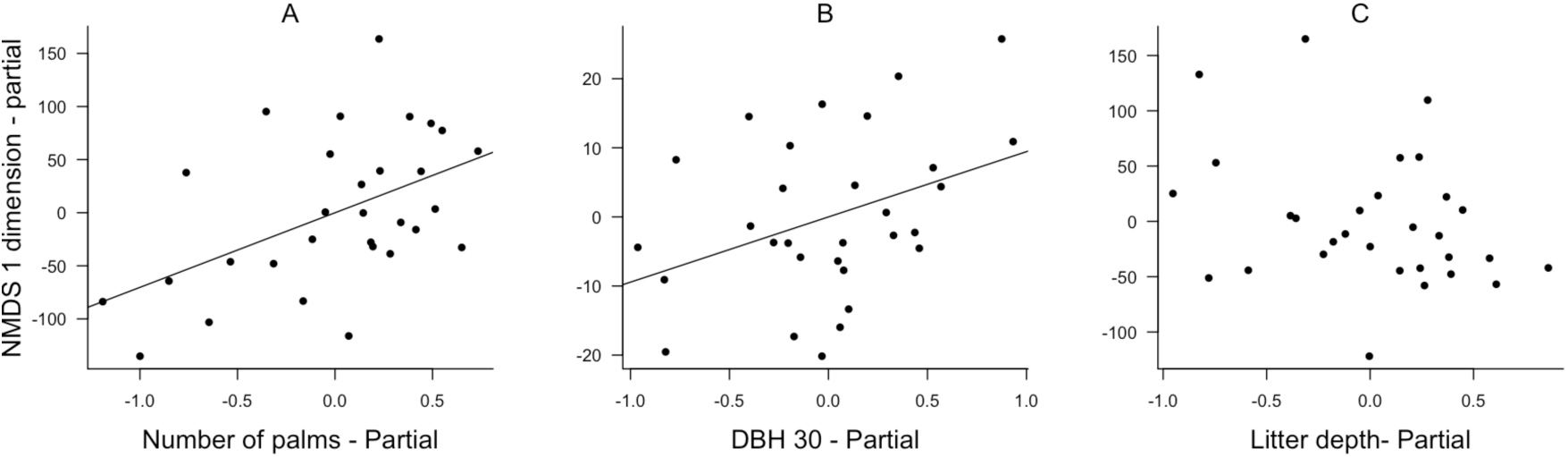
Partial regressions of the significant effects detected on species composition (NMDS Axis) of harvestmen. a) Species composition change with the increment of number of palms (P = 0.005), and b) number of tress with diameter at breast high > 30 cm (P = 0.034), and c) species composition was not affected by litter depth (P > 0.1).

**Table 3.** Multiple regression models for species composition (NMDS Axis), richness and abundance

| **Response** | **Predictor** | **Coefficient** | **Standard Error** | **t** | **P** | **model R<sup>2</sup>** |
| --- | --- | --- | --- | --- | --- | --- |
| NMDS Axis | Intercept | -2.11 | 0.763 | -2.766 | 0.01 | 0.417 |
|  | Npalms | 0.003 | 0.001 | 3.064 | **0.005** |  |
|  | DBH30 | 0.015 | 0.007 | 2.164 | **0.039** |  |
|  | Litter | -0.002 | 0.001 | -1.631 | 0.114 |  |
| Richness | Intercept | 10.036 | 4.654 | 2.157 | 0.04 | 0.014 |
|  | Npalms | -0.002 | 0.007 | -0.39 | 0.699 |  |
|  | DBH30 | -0.001 | 0.044 | -0.264 | 0.794 |  |
|  | Litter | -0.001 | 0.008 | -0.055 | 0.956 |  |
| Abundance | Intercept | 64.141 | 15.51 | 4.135 | 0.001 | 0.245 |
|  | Npalms | -0.036 | 0.024 | -1.502 | 0.145 |  |
|  | DBH30 | -0.21 | 0.147 | -1.428 | 0.165 |  |
|  | Litter | -0.039 | 0.028 | -1.389 | 0.176 |  |

## Discussion

Harvestman diversity was related to local and meso scale gradients created by vegetation structure at this forest. At local scale, trees harbored the most diversity and abundant harvestman assemblages, compared with other available habitats. At meso scale number of trees and palms were the best predictors of harvestman composition, while both were not related with harvestman richness or abundance. In context, the harvestmen diversity at Reserva Ducke is comparable with recent harvestmen inventories carried out in different localities of *upland* forest across the Amazon region (Pinto-da-Rocha & Bonaldo, 2006; Bonaldo *et al*., 2009; Tourinho *et al*., 2014, Porto *et al*., 2016), with number of collected species ranging between 26 to 30. Cosmetidae and Sclerosomatidae were the most abundant families, while Stygnidae and Sclerosomatidae were the most diverse groups, with five and four species collected each.

### Harvestmen diversity at local scale

Large trees can offer microhabitats such as cracks, bark pockets, bark pockets with decay, bowls in bark, stem cavities, witch broom, hollow chambers on butt of trees, among other 12 tree microhabitats defined by Michel and Winter (2009), which are probably suitable for harvestmen. A total of 23 species out of 27 were found foraging on trees, indicating that trees might be one of the most important habitats for harvestmen at local scale. From these, six species were found exclusively on trees. These species were observed mainly on trees with a highly corrugated bark and big roots (locally known as *sapopemas*), suggesting that harvestmen could be using bark pockets, cracks and cavities as refuge or for prey source. For instance, most individuals of Samoidae Genus 1 sp. 1 were found while foraging in tree bark pockets and cracks.

The use of tree bark may be related to harvestmen size. With the exception of the gonyleptids of the genus *Discocyrtus*, all species found only on trees were small (dorsal scutum length < 2.5 mm). In addition, due to their larger mass, large diameter trees provide temperature-buffering microhabitats (Brower *et al*., 2009). This could be of benefit for some harvestmen species, especially the smaller ones that probably are more vulnerable to changes in temperature and moisture.

Abundance of harvestman per tree was also higher in larger trees than in smaller trees or shrubs for this assemblage. This can also be related to quantity of microhabitats available in the tree bark. In addition, large harvestman species can also take advantage of trees, as known for the cranaid *Phareicranaus manauara*, which uses trees in its reproductive strategy (Colmenares & Tourinho, 2014).

Despite the high aggregation on trees, some harvestman species were sampled in different habitats. Species of Sclerosomatidae were more generalist, occupying almost all microhabitats, with *Caluga* sp. 1 as the only species distributed across all available local habitats. The longer lengths of legs and the high number of tarsomeres, which increase their capacity to climb and reach upper places on the vegetation, can explain this observation (Adams, 1984; Proud *et al*., 2012). However, *Prionostemma* sp. is probabily specialized in using some specific mature tree trunks that have surface color patterns similar to its body color, facilitating camuflage. From the 34 individuals collected, 32 were on trees while only 2 were found in bushes and leaf litter.

### Relationship between vegetation structure and harvestmen assemblages

A more complex habitat can allow the co-occurrence of more harvestmen species by increasing the availability of microhabitats (Proud *et al*., 2012). It is known that diversity and quantity of microhabitats increases with tree diameter, promoting their use by vertebrates and invertebrates and acting as predictor of biodiversity (Michel & Winter, 2009). Thus, more trees with DBH above 30 cm per plot should mean more microhabitats available for all invertebrates in the study area, including harvestmen. However, our data show that either species richness or abundance at the meso scale may not be directly related with number of trees or palms. However, species composition per plot was related with vegetation structure predictors. Thus, plots with a higher number of large trees or higher number of palms may not affect the total number of species or individuals, but harbors different assemblages composition. Palm dwellers, can show lower abundances in plots with less palm trees, while trees dwellers, can be more abundant in plots with more large trees available.

We observed that harvestman use both faces of palm leaves for foraging, and Vasconcelos *et al*., (1990) suggested that acaulescent palms can increase habitat complexity, as a consequence of the fallen litter trapped on their leaves. Tourinho *et al*., (2014) and Porto *et al*., (2016) also suggested that palms might be reflecting the overall variation of habitat structure. Thus, we can hypothesize that, at least for Reserva Ducke, more palms in a given plot would proportionally change the availability of other kinds of microhabitats, such as the ones related to trees, dead trunks and leaf-litter trapped in acaulescent palms. Consequently, plots with higher number of palms would proportionally harbor more palm dwellers species. The same relation may be applied to number of trees per plot.

Harvestman species composition can also be affected by the increment in the numbers and abundance of generalist and vegetation dwellers, which are usually more abundant than tree and ground dwellers. For instance, *Caluga* sp. 1, *E. duapunctata*, *Prionostemma* sp. 1 and *Protimesius longipalpis*, among others, could benefit from the number of palms, but species like Samoidae Genus 1 sp. 1, Gonyleptidae spp., and Zalmoxidae *Chamaia* sp. 1, more related to trees, can be limited by the decrease of adequate microhabitats. Moreover, two of the recorded species use palms in their reproductive strategies. The stygnid *P. longipalpis* was recorded during our fieldwork using fallen palm trunks as an alternative refuge for the nymphs. In Stygnidae, at least another two species have been seen using palms leaves as a substrate to keep their clutches (Villarreal & Machado, 2011). There is also evidence that *P. manauara* and other *Phareicranaus* use fallen palm leaves and trunks to place their clutches and/or keep their nymphs (Hunter *et al*., 2007; Proud *et al*., 2011; Colmenares & Tourinho, 2014).

It is widely known that large old trees sustain countless other species, their hollows and crevices shelter many different animals and their branches and trunks are real diverse gardens (Lindenmayer & Laurance, 2016). However, they are susceptible to several threats including deforestation, logging, agriculture, drought, fire, windstorms, invasive species, the development of human infrastructure, and climate change. Across the planet old growth-forest have been cleared for human use and in the Amazon the mortality rates of large old trees are growing very fast (Lindenmayer *et al*., 2012). Our data suggest that areas of *upland* forest in the central Amazon with a balance between number of large trees and palms are suitable to maintain a comprehensive assemblage of harvestman species. Consequently, any disturbance resulting in reduction of the number of large trees will have a strong impact on harvestmen diversity, especially by limiting the occurrence of tree dwellers species. We know very little about the relationship of other non-flying arthropod and tree structure, however, our results indicate the conservation of large old trees and their global decline must be taken into consideration as a major concern to keep harvestman diversity in Amazon rainforest.

## Acknowledgements

José W. de Morais (INPA) helped with the transportation for the fieldwork. Isadora Williams and Willians Porto helped as field assistants. Gonzalo Giribet kindly helped with the identification of the Cyphophtalmi. The material for this study was obtained under collecting permits 39557, granted from SISBIO to the first author. This study was supported by the Coordination of Improvement of Higher Education Personnel - CAPES (PEC-PG grant #5828104 to PAC and PNPD grant #03017/09–5 to ALT), Brazilian National Council for Scientific and Technological Development – CNPq through the Brazilian Research program for Biodiversity – PPBio, National Institute of Science and Technology for Amazonian Biodiversity (INCT-CENBAM), The Program Science without borders Special Visiting Professor (CSF/PVE) and The Program Science without borders for both the Special Visiting Professor Grant (CAPES #003/2012), International Postdoctoral grant (CNPq #200972/2013–8 to ALT) and Foundation Lemann for the Lemann additional International fellowship to ALT.

